# GENOME-WIDE IDENTIFICATION, CHARACTERIZATION AND EXPRESSION ANALYSIS OF NA^+^/H^+^ EXCHANGER GENE FAMILY IN TWO CORCHORUS SPECIES

**DOI:** 10.1101/2024.04.01.587583

**Authors:** Borhan Ahmed, Anika Tabassum, Kazi Khayrul Bashar, Nasima Aktar, Sabbir Hossain, Abdullah-Al Mamun, Shyduzzaman Roni, Fakhrul Hasan, Mobashwer Alam

**Affiliations:** Bangladesh Jute Research Institute, Manik Mia Avenue, Dhaka-1207; American International University of Bangladesh, Dhaka; Bangladesh Forest Research Institute, Chittagong, Bangladesh; Faculty of Agriculture, Bangabandhu Sheikh Mujibur Rahman Agricultural University, Gazipur, Bangladesh; The University of Queensland

## Abstract

Na^+^/H^+^ exchangers (NHXs) are secondary ion transporter to exchange H+ and transfer Na^+^ or K^+^ across membrane plays crucial role in cellular pH and Na^+^ and K^+^ homeostasis in plant. In the present study, we have performed genomewide identification, characterization and comparative analysis of NHXs genes in two cultivated jute species and found seven Co and Cc NHXs in each genome. Subsequently phylogenetic analysis demonstrated that Co and Cc NHXs have three sub-group that correlate with other plant species containing 10-12 transmembrane helices (TMH) and a hydrophilic C-terminal domain in the cytoplasmic region, the TMH compose a hollow cylinder to provide the channel for Na^+^ and H^+^ transport. Promoter profiling of both species NHXs genes revealed in enrichment of cis element involved in stress responsiveness reflecting their potential involvement against abiotic stress. Transcriptome data and real time PCR of Co and Cc NHXs gene expression analysis under saline and drought condition showed significant differences, which suggest their potential role in abiotic stress conditions. This is the first comprehensive analysis providing baseline information of Co and Cc NHXs genes in abiotic stress condition which can be used for the development of abiotic stress tolerant jute variety.

## Introduction

The salinization of soil has become a widespread environmental problem limiting agricultural productivity worldwide. High salinity leads detrimental effects through three main aspect, initially saline soil conduct osmotic stress by inhibiting the plant normal water uptake. Then salt stress induces Na^+^ concentration in the cytosol of plants which are toxic to important physiological and biochemical process^1,2^. Finally high soil salt concentration can also induce oxidative stress, which can cause a series of oxidative damage^3^. In saline tolerant plant, they have evolved some special mechanisms to minimize the accumulation of toxic ions in plant tissue by partitioning them in the apoplast and vacuole, increases the synthesis of osmotic adjustment substances such as proline and betaine for maintaining tissue water status, and enhance antioxidant capacity to prevent the occurrence of oxidative stress^4^. Plant cell have two strategies to maintain a high concentration of K^+^ and a low concentration of Na^+^ in the cytosol, Na^+^ efflux and vacuolar compartmentalization of Na^+^. Na^+^ efflux is dependent on a plasma membrane Na^+^/H^+^ exchanger (NHX) that is encoded by the salt overly sensitive 1 (SOS1) gene in Arabidopsis^5^. By contrast, vacuolar NHXs catalyze the compartmentalization of Na^+^ into vacuoles, which is another important strategy for salt tolerance in plants. The sequestration of Na^+^ in vacuoles maintains a lower Na^+^ concentration in cytoplasm, which relieves the toxic effect on cytosolic enzymes and lowers the osmotic potential in the vacuole, maintaining turgor pressure and cell expansion under salt stress conditions^1,6^.

To date, all sequenced eukaryotes containing multiple ubiquitous NHX genes, except for yeast contain a single NHX^7^. Based on their subcellular localization NHXs are classified into three major classes: Plasma membrane (PM) class, endosomal (Endo) class and vacuole (Vac) class^8^. In Arabidopsis, NHX are consisting of six intracellular members belongs to Vac-class (AtNHX1-4) or in Endo-class (AtNHX5-6), and two divergent member located at the plasma membrane^7,9^. Briefly, intracellular members (AtNHX1-6) transport either Na^+^ or K^+^ into the vacuoles or endosomal region in exchange for H^+^ efflux to the cytosol, while plasma membrane-bound members (AtNHX7-8) transport Na^+^ out of the cell in exchange for H^+^ influx into the cell^9,10^. Plant NHXs mediate both Na^+^/H^+^ and K^+^/H^+^ exchanges^11-13^, therefore, they affect both salinity tolerance and K^+^ nutrition. Many studies have suggested that NHX overexpression leads to improved salt tolerance in diverse species^14-20^.

NHXs involve in various biological processes such as salt stress response^21,22^, pH homeostasis^23,24^, K^+^ homeostasis^12,25^, cell expansion^11,22^, cellular vesicle trafficking^26,27^. In Arabidopsis, the double knockout nhx1 and nhx2 had significantly reduced growth, smaller cells, shorter hypocotyls in etiolated seedlings and abnormal stamens in mature flowers as compared with the single knockouts nhx1 and nhx2^28^. Furthermore, the double knockout nhx5 nhx6 missorted vacuolar destined cargo to the apoplast, and the Golgi and trans-Golgi network in nhx5 nhx6 were significantly more acidic than in wild type^22^ suggesting that endosomal NHXs have important roles in the ion trafficking of plant cell by regulating vesicular pH homeostasis^29^.

Before the year 2000, limited knowledge was available of how abiotic stresses induces NHX genes, despite recognizing the role of NHXs as enzyme involved in salinity tolerance. To date, the NHXs have been identified in the genome of numerous plants, including 8 in *A. thaliana*^30^, 7 in rice^31^, sorghum^32^, maize^33^, *Medicago truncatula*^34^, 8 in grape^35^ and Poplar^36^.

We have identified 7 putative non-redundant NHX gene in both jute species by Insilico cloning. Nomenclature and classification were performed based on phylogenetic analysis according to the model plant Arabidopsis. To provide holistic comprehensive insights into NHX members in two Corchorus species, we analyzed exon-intron junctions as well as putative conserve residues involved in substrate specificity, with a focus on sub-cellular localization and prediction of transmembrane domains. We also profiled gene expression of C capsularis and C. olitorius NHX genes in response to salt and drought stress.

## MATERIALS AND METHODS

### Genomic data mining and NHX gene identification

To identify the NHX protein sequence in *C. capsularis* and *C. olitorius*, reference proteins of well-established NHX protein sequence were chosen as query sequences. The validated reference proteins of Arabidopsis were downloaded from TAIR database (http://www.arabidopsis.org); Poplar from phytozome (http://phytozome.jgi.doe.gov/), NCBI database (http://www.ncbi.nlm.nih.gov/) and both Corchorus sp. Genome sequence from barj database. The downloaded sequences were used as query to perform blast with the both jute species genome database and identified the putative homolog in jute species. The BLAST server National Center for Biotechnology Information (http://www.ncbi.nlm.nih.gov/BLAST/) was used for identification with a cut off e-value of e-10. The amino acid sequence of candidate genes was analyzed to examine the presence of the characteristic Conserved domains within the Co and CcNHX protein sequences using NCBI’s Conserved Domain Database (CDD, www.ncbi.nlm.nih.gov/Structure/cdd/cdd.shtml) SMART (http://smart.embl-heidelberg.de/)^37^, Pfam (http://pfam.xfam.org/)^37^ and InterProScan (http://www.ebi.ac.uk/tools/pfa/iprscan5/)^38^ analysis to verify their presence. All the identified genes were manually checked, and the predicted genes without having the identical domain, were rejected. Transmembrane helical domains (TMHs) were assessed by TMHMM Sever v.2.0 (http://www.cbs.dtu.dk/services/TMHMM/) and SOSUI (http://harrier.nagahama-ibio.ac.jp/ sosui/sosui_submit.html); the subcellular localization of Co and CcNHXs were predicted using PlantmPLoc (http://www.csbio.sjtu.edu.cn/bioinf/plant-multi/). The results were then manually curated for altered and missing transmembrane domains.

### Gene structure analysis

In silico approaches were used to obtain the genomic sequences of the identified NHX gene family. Exons and introns structures diagram of NHX genes in both jute genomes were determined by a web application named Gene Structure Display Server using GSDS^39^ (http://gsds.cbi.pku.edu.cn/).

### Motif analysis NHX family proteins

Genome-wide and molecular evolution analyses of the NHX gene family was performed in both jute genome using Multiple expectation maximization for motif elicitation (MEME) utility program according to the method described by Bailey et al. (2009)^40^.

### Physico-chemical characterization of NHX gene family proteins

Assessment of physical and chemical properties of the NHX gene family proteins was done with the help of ProtParam online (http://expasy.org/tools/protparam.html) and related indexes, including theoretical isoelectric point (pI), molecular weight, formula, aliphatic index, instability index, and grand average of hydropathicity (GRAVY).

### Phylogenetic analysis of NHX gene family

To gain more insight into the evolution of the NHX gene family in both Corchorus genome, a total of 7 genes were identified from both Corchorus species. Multiple sequence alignments of the amino acid sequence of NHX genes of both species along with *Arabidopsis thaliana* were generated using clustalW with the default settings. The bootstrap consensus phylogenetic tree inferred from 1000 replicates was constructed using the maximum likelihood (ML) method with MEGA6^41^.

### Promoter regions analysis of NHX gene family genes

To investigate cis-elements in promoter sequences of NHX family genes, 1000bp of both Corchorus species genomic DNA sequences upstream intergenic region from the initiation codon (ATG) were extracted from jute genome database. And then the PlantCARE (http://bioinformatics.psb.ugent.be/webtools/plantcare/html/)^42^ and PLACE (Higo et al., 1999) (http://www.dna.affrc.go.jp/PLACE/)^43^ was used to analyze cis-elements in the promoters.

### Mapping both Corchorus species NHX genes on chromosomes

The physical location of Co and Cc NHXs of each Corchorus species chromosome/scaffold was detected using BLASTNT search against the local database of the both jute genome. Starting position of all Co and Cc NHXs genes were used as the indicative position of the genes on the chromosome or the scaffold. Tandem duplications of NHX genes in the both species genome were identified by checking their physical locations within 100-kb adjacent region in individual chromosomes^44^. Due to the unavailability of the full chromosome-scale assembly is both jute genome, the assembled sequence in the form of scaffolds was helpful to locate tandem duplications.

### Plant materials

O-4, an important cultivar in Bangladesh of C. olitorius and CVL1 an widely cultivated cultivar of C. capsularis were hydroponically cultivated at 25-28°C in a green house. Each genotype was grown in two container containing 120 plant per container. Uniformly germinated seeds were grown in ½ strength Hoagland nutrient solution^45^, which was replaced with fresh solution every third day. After 25 days (nine leaf stage), saline condition was created by supplementing Hoagland solution with 250 mM NaCl and plants at Hoagland solution was used as control. After 0, 2, 4, 8h of saline treatment, uniformly growth jute leaf and root sample were cut off with a sharp scalpel. A total of 24 samples from four different time point were carefully harvested, immediately frozen in liquid nitrogen and store at -80°C for RNA extraction.

### Total RNA extraction and cDNA synthesis

One gram of tissue samples was disrupted in liquid nitrogen to a fine powder using a mortar and pestle. Total RNA was extracted from all samples using an in-house modified Cetyltrimethyl ammonium bromide (CTAB) protocol (unpublished results). Total RNA concentration and purity were determined using a NanoDrop 2000 spectrophotometer (NanoDrop, Thermo Scientific, USA). The integrity of total RNA was evaluated by 1% agarose gel electrophoresis. The absence of genomic DNA in the sample was confirmed by PCR using crude RNA without reverse transcription as a template and assessed by 1.2% agarose gel electrophoresis. Before cDNA synthesis, total RNA was treated with amplification grade DNase I (Sigma Aldrich, Germany) to remove any traces of genomic DNA according to the manufacturer’s instructions. Subsequently, the first strand complementary DNA (cDNA) was synthesized from 2 µg of total RNA using the RevertAid First Strand cDNA Synthesis Kit (Thermo Fisher Scientific, USA) according to the manufacturer’s instructions. Following these steps, the samples were incubated with RNaseH (Thermo Fisher Scientific, USA) to degrade the RNA strand of any RNA-DNA hybrids according to the manufacturer’s instructions. The cDNA was then stored at -20 °C.

### Primer design and amplification efficiency

All obtained NHX coding sequences were aligned (Fig.1) and divided into three different groups based on phylogeny and visual interpretation of multiple sequence alignment and sequence similarities in the untranslated region. Primer pairs for the NHX genes (7+7 in two Corchorus species) were designed using an online tool from IDT (http://sg.idtdna.com/) and GenScript (www.genscript.com). When possible, the primers were designed to span an exon-exon junction to avoid any genomic DNA contamination (Supplementary table 1). The specificity of these primer pairs was evaluated by PCR followed by 1.2% agarose gel electrophoresis and melt curve analysis during qRT-PCR. Calculation of the amplification efficiency was based on the slope of the standard curve. Standard curves were generated by performing qRT-PCR for each gene using serially diluted cDNA samples. The amplification efficiency was then calculated using the following formula: E = (10 -1 /slope-1) × 100%.

**Figure 1.**
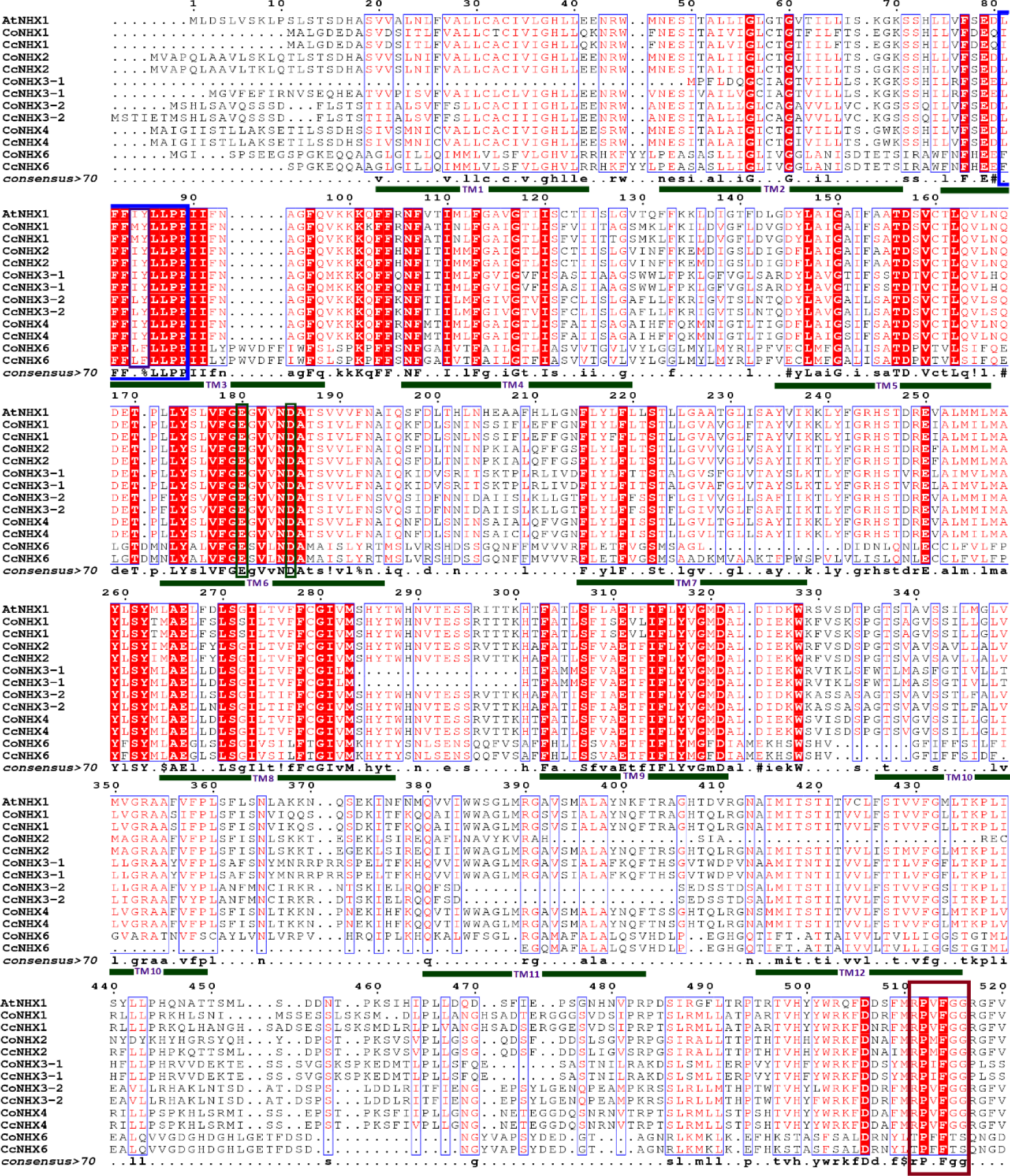
Alignment of amino acid sequences of the Co and CcNHX family genes. Protein sequence were aligned using Clustal Omega software. Putative transmembrane domains are represented by black lines over the sequence using Roman numerals. The amiloride binding motif is represented by a maroon colour in third transmembrane domain. The residues important for antiporter activity are marked by dark green colour. TheC-terminal region important for antiporter activity is marked by a red box. The marking among the sequence is done by Jalview software^55^.

### qRT-PCR expression analysis

Changes in the expression of Co and Cc NHX genes in leaf and root of different time stage in response to salt stress were detected by quantitative real-time polymerase chain reaction (qRT-PCR) analysis using LightCycler® 480 SYBR Green I Master mix (Roche Diagnostics, Germany) chemistry on a Real Time PCR System (Applied Biosystems) instrument. The 10µl reaction volume consisted of the following: 5 µl SYBR mix, 1 µl (10pmol) each forward (F) and reverse (R) gene specific primer, 1 µl template cDNA (50ng) and 2 µl PCR water. The conditions for real-time PCR were as follows: initial denaturation at 95°C for 5 min, followed by 40 cycles of denaturation at 95°C for 10s, annealing at 58°C for 10s, and extension at 72°C for 15s. The qRT-PCR reactions were normalized using the house keeping gene Catalytic subunit of protein phosphatase 2A (PP2A) and ubiquitin conjugating enzyme E2 (UBC2), reference for all comparisons (unpublished result). The florescence was measured following the last step of each cycle and three replications were used for each sample. The relative expression level of the target genes were assessed based on 2^-^^CT^ methods^46^.

### Expression Analysis

Publicly available jute transcriptome (RNA-Seq) data against drought and salinity stress along with seedling and fiber cell transcriptome data were used for the study of expression pattern of NHX gene family in two jute species and were downloaded from NCBI sequence Read Archive (SRA) database. Both Corchorus olitorius (variety name GF) and Corchorus capsularis (variety name YY) control and drought stressed data were downloaded from SRA (accession id: SRR6429829, SRR6429828, SRR6429827, SRR6429826, SRR6429825, SRR6429824, SRR6429823, SRR6429822, SRR6429821, SRR6429820, SRR6429819, SRR6429818)^47^. Similarly, to determine the effectiveness of NHXs genes against salinity the RNA-Seq data from leaf and root of one capsularis variety (YY) and two olitorius variety (Salt sensitive NY and salt tolerant TC) from SRA database were downloaded through the following accession id: SRR6308432, SRR6308433, SRR6308434, SRR6308435, SRR6308436, SRR6308437, SRR6308438, SRR6308439, SRR6308440, SRR6308441, SRR6308442, SRR6308443, SRR6308368, SRR6308369, SRR6308370, SRR6308371, SRR6308372, SRR6308373, SRR6308374, SRR6308375, SRR6308376, SRR6308377, SRR6308378, SRR6308379)^48,49^. High quality clean RNA-Seq reads were mapped to the both Corchorus species data by Tophat2 (version 2.1.0, Baltimore, MD, USA) with the default settings^50^. Quantification of gene expression was done using the Fragments Per Kilobase Of Exon Per Million Fragments Mapped (FPKM) with Cuffdiff algorithm^51^. The clustered heatmap of Z scaled resulting FPKM values of Co and Cc NHXs was generated using the heatmap function of R package (version 3.2.2; available online: https://cran.r-project.org/web/packages/pheatmap/).

## Result

### Identification and physico-chemical properties of both species NHX gene family

BLAST analysis of NHX genes in Arabidopsis and polar resulted a total of 7 putative NHX encoding genes in both jute genome, named thereafter as CoNHXs and CcNHXs (Supplementary table1). The Na+/H+ exchanger domain (PF00999) predicted through the online program SMART (http://smart.embl-heidelberg.de/)^37^ with default e-value for further confirmation. The number of NHX genes are almost similar with most of the species named *Oryza sativa*, *Medicago truncatula*, *Sorghum bicolor*, *Zea mays* and one number lower than *Arabidopsis thaliana*, *Populus trichocarpa*, *Vitis vinifera*^36^. The sequences of NHX genes between two Corchorus species are almost same. The information analysis of predicted sequence of both Corchorus species NHXs ranged from 412-543 amino acid with average 511aa (C. olitorius 461-543aa and C. capsularis 412-543aa) (supplementary table 2) in vacuolar and endosomal NHXs where as protein length of plasma membrane NHX7/SOS1 is more than double compare to vacuolar and endosomal NHX (1152 and 1159 aa in C. olitorius and C capsularis respectively). Similar features observed in molecular weight of NHX genes, where Vacuolar and endosomal NHX protein are average 56.46 kDa and plasma membrane NHX protein is 128.29 kDa. According to phylogenetic tree there are three different groups of NHX gene family and the first is vacuolar NHX (NHX1-4) group is alkaline in nature (PI: 7.23-9) but second and third gene groups (NHX5-7) are acidic (PI: 5.14-6.15) in nature. The GRAVY value of a protein is calculated as the sum of the total number of residues present in the sequences. Positive and negative GRAVY scores reflect hydrophobicity and hydrophilicity respectively. The GRAVY scores of both Corchorus species NHXs were positive which indicated that all were hydrophobic proteins. However the degree of hydrophobicity exhibited higher in vacuolar and endosomal NHXs (0.402 to 0.628) but very low in plasma membrane NHX (NHX7/SOS1) which is 0.047 in C. capsularis and 0.072 in C. olitorius.

### Subcellular localization analyses

Based on subcellular localization to vacuole, endosomal compartment and plasma membrane NHX protein are classified into three major classes, namely, Class I, Class II and Class III^8^. As there is no genome-specific subcellular localization prediction software available for Corchorus species, we performed a bioinformatics analysis using a generalized plant localization predictor, Plant-mPLoc^52^. The localization of the class II protein to endosomal compartments was confirmed based on its transmembrane domains^53^. Based on the predication, Co and CC NHX1 to 5 were localized to vacuole, Co and Cc 6 was localized to Golgi, and Co and CcNHX7 was predicted to be localized to plasma membranes (Table 1) which were in line with the localization of their corresponding AtNHX proteins^8^.

**Table 1.** Physico-chemical properties of NHX gene family in two Corchorus species.

| SI No | Gene Name | Protein length | Theoretical pI | Molecular Formula (Da) | Aliphatic index | Instability index | GRAVY | Molecular weight | Subcellular localization |
| --- | --- | --- | --- | --- | --- | --- | --- | --- | --- |
| **Corchorus olitorius NHX genes** |  |  |  |  |  |  |  |  |  |
| 1 | CoNHX1 | 527 | 8.93 | C<sub>2689</sub>H<sub>4206</sub>N<sub>664</sub>O<sub>733</sub>S<sub>19</sub> | 110.44 | 37.12 | 0.518 | 58174.1 | Vacuole |
| 2 | CoNHX2 | 502 | 7.59 | C<sub>2570</sub>H<sub>3950</sub>N<sub>634</sub>O<sub>702</sub>S<sub>21</sub> | 104.66 | 42.48 | 0.462 | 55634.7 | Vacuole |
| 3 | CoNHX3 | 461 | 9.00 | C<sub>2423</sub>H<sub>3743</sub>N<sub>579</sub>O<sub>633</sub>S<sub>21</sub> | 109.09 | 39.42 | 0.574 | 51786.1 | Vacuole |
| 4 | CoNHX4 | 543 | 8.15 | C<sub>2748</sub>H<sub>4291</sub>N<sub>683</sub>O<sub>756</sub>S<sub>25</sub> | 109.13 | 37.79 | 0.518 | 59794.9 | Vacuole |
| 5 | CoNHX5 | 517 | 6.15 | C<sub>2618</sub>H<sub>4091</sub>N<sub>639</sub>O<sub>735</sub>S<sub>21</sub> | 110.87 | 41.26 | 0.572 | 56951.3 | Vacuole |
| 6 | CoNHX6 | 516 | 5.26 | C<sub>2632</sub>H<sub>3994</sub>N<sub>644</sub>O<sub>728</sub>S<sub>23</sub> | 100.50 | 43.41 | 0.430 | 57043.9 | Endosomal |
| 7 | CoNHX7/SOS1 | 1152 | 6.16 | C<sub>5789</sub>H<sub>9140</sub>N<sub>1526</sub>O<sub>1669</sub>S<sub>35</sub> | 104.77 | 35.73 | 0.072 | 127943.5 | Plasma membrane |
| **Corchorus capsularis NHX genes** |  |  |  |  |  |  |  |  |  |
| 1 | CcNHX1 | 532 | 8.93 | C<sub>2711</sub>H<sub>4238</sub>N<sub>676</sub>O<sub>741</sub>S<sub>18</sub> | 109.59 | 35.73 | 0.482 | 58734.6 | Vacuole |
| 2 | CcNHX2 | 543 | 7.23 | C<sub>2758</sub>H<sub>4302</sub>N<sub>678</sub>O<sub>748</sub>S<sub>26</sub> | 110.04 | 36.12 | 0.584 | 59760.1 | Vacuole |
| 3 | CcNHX3 | 508 | 8.42 | C<sub>2656</sub>H<sub>4124</sub>N<sub>640</sub>O<sub>697</sub>S<sub>23</sub> | 113.94 | 39.61 | 0.628 | 56911.2 | Vacuole |
| 4 | CcNHX4 | 543 | 8.11 | C<sub>2748</sub>H<sub>4290</sub>N<sub>684</sub>O<sub>755</sub>S<sub>26</sub> | 108.95 | 37.77 | 0.518 | 59823.9 | Vacuole |
| 5 | CcNHX5 | 523 | 5.74 | C<sub>2648</sub>H<sub>4124</sub>N<sub>642</sub>O<sub>747</sub>S<sub>22</sub> | 108.85 | 42.24 | 0.567 | 57611.0 | Vacuole |
| 6 | CcNHX6 | 412 | 5.14 | C<sub>2079</sub>H<sub>3154</sub>N<sub>508</sub>O<sub>593</sub>S<sub>20</sub> | 94.85 | 38.95 | 0.402 | 45394.1 | Endosomal |
| 7 | CcNHX7/SOS1 | 1159 | 6.20 | C<sub>5814</sub>H<sub>9188</sub>N<sub>1536</sub>O<sub>1682</sub>S<sub>35</sub> | 103.72 | 35.06 | 0.047 | 128640.2 | Plasma membrane |

### Transmembrane domain analysis

The transmembrane domain were detected using TMHMM and SOSUI software tools and predicted 12 transmembrane domain (Supplementary table2-3) along with a putative Amiloride-binding domain in TM3 in both Corchorus species NHXs genes (Fig. 1). A putative cation-binding domain exists between TM5 and TM6, and this region shared a high similarity with the corresponding cation-binding domain ofAtNHX1. This region of AtNHX1 has been postulated to contain the putative cation-binding domain as mapped by homology to other exchangers^21^. The alignment shows that the C terminal ends of Co and Cc NHXs are less conserved compared with the remaining parts of the protein sequences (Fig. 1). However, Co and Cc NHX1-5 have a highly conserved region in the end of their C-termini, which shares a high similarity with the calmodulin (CaM)-like protein 15-binding domain of AtNHX1^54^.

### Phylogenetic analysis

To explore the evolutionary relationship among the NHX gene family members were studied by computing an unrooted phylogram showing divergence in Arabidopsis thaliana, Corchorus olitorius and Corchorus capsularis. External nodes in the phylogram represent each protein sequence, and internal nodes display the percentage support for each divergence. Branch distances in the tree are shown in units of residue substitutions per site. The phylogenetic analysis revealed that each individual class in NHX falls under a separate clade. Co and CcNHX1 through 5 (class I) were in one clade, Co and CcNHX6 (class II) in another clade, and Co and Cc NHX7 (class III) in a separate clade (Fig.2). AtNHX5 and AtNHX6 fell in the same clade as Co and CcNHX6, and AtNHX7 and AtNHX8 fell in the same clade as Co and CcNHX7 (Fig.2). AtNHX1 and AtNHX2 were very similar; however, corresponding genes in C. olitorius and C. capsularis separated into different groups (Fig.2).Based on the evolutionary distance, Co and CcNHX4 appeared to be more closely associated with Co and CcNHX1 and 2.

**Figure 2.**
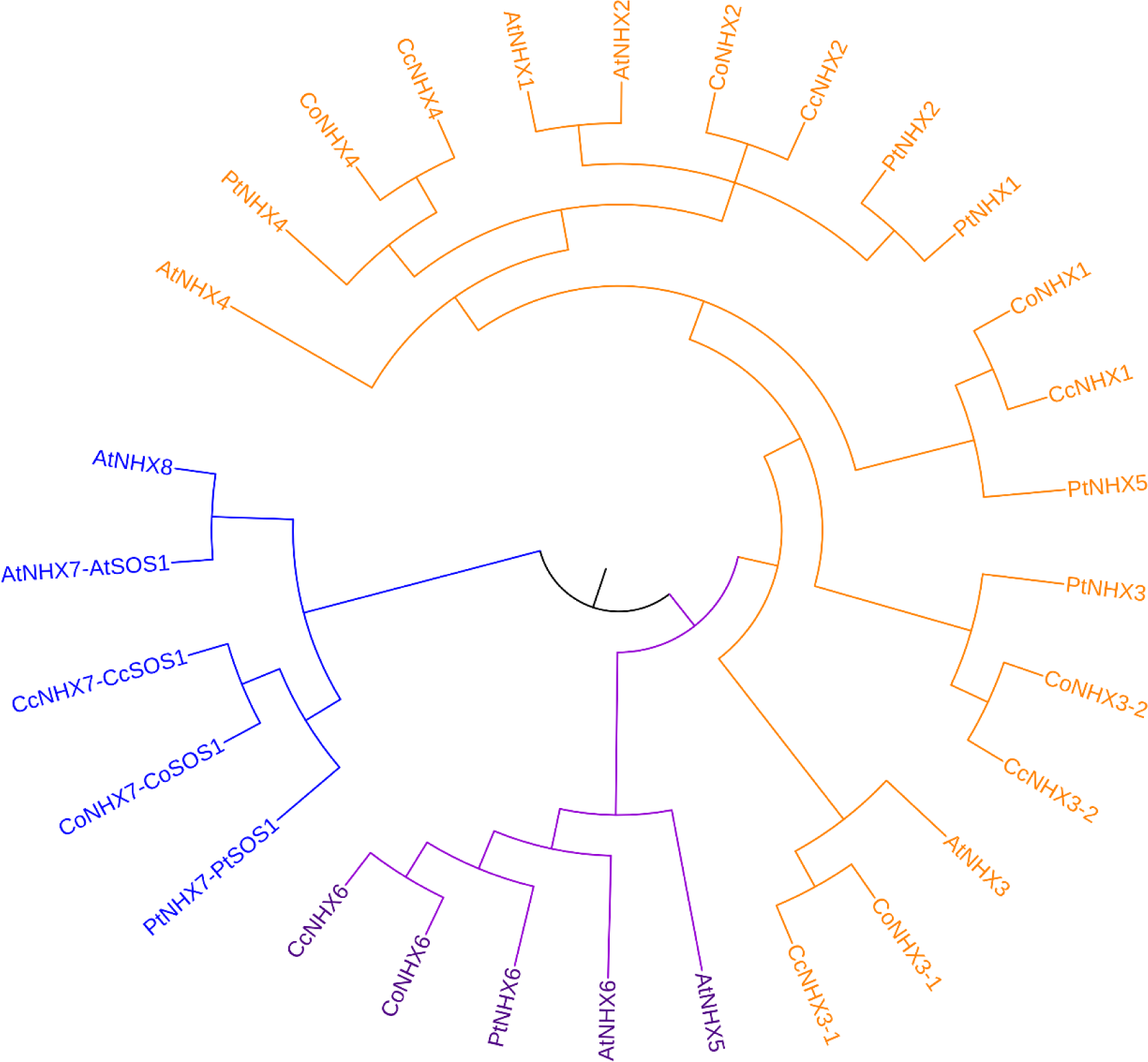
Phylogenetic analysis of the NHX genes in A. thaliana, C. olitorius, C. capsularis. Phylogenetic was constructed using maximum likelihood (ML) method with 1000 bootstrap replicates. The three major classes are marked with different colour.

### Exon intron organization

To analyze the structural characteristics of the two *Corchorus species NHX* genes, we compared the exon-intron organizations of these genes (Fig. 3). Among the Co and CcNHX genes, Vac-class NHXs (Co and Cc NHX1-4) had 13 introns and Co and CcNHX5 had 12 intron, In endo-class 20 and 17 intron observed in CoNHX6 and CcNHX6 respectively, While PM-class *NHXs* (Co and CcNHX7) had 22 introns. The genomic organization of Co and CcNHX7 contain long gap with in first intron and CoNHX6 contain largest 9th intron and CcNHX6 hold long intron in 8th position. The structures and sizes of the exons were found to be well conserved among Co and CcMnNHX1-5. However, the exon-intron structure of Co and CcNHX6 and NHX7 were significantly different from that of Co and CcNHX1-5. These results were corroborated by the phylogenetic analysis, indicating that the both Corchorus species NHX family genes form three subgroups.

**Figure 3.**
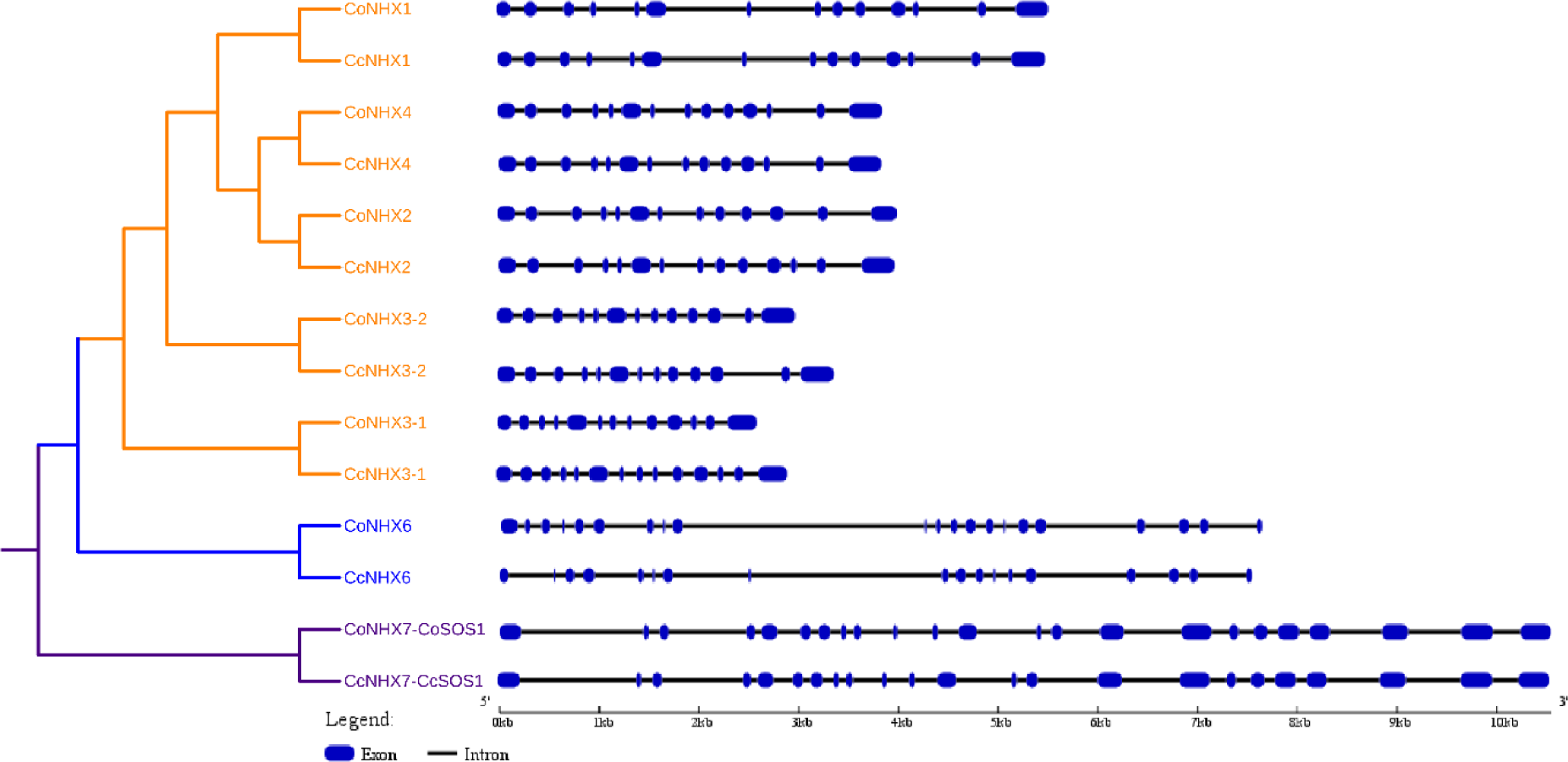
Structural organization of Co and CcNHX genes. Exon are represented as rectangles, intron are represented as lines. Structural organization was generated using Gene Structure Display Server.

### Analysis conserved motifs

To further investigate the characteristic region of Co and CcNHX proteins, the motif distributions in Co and CcNHX proteins were analyzed using MEME and total of 10 individual motifs were identified (Fig. 4 and Supplementary table4 and SupplementaryFig.1). As predicted, the closely related members had common motif composition. Six motifs (motif 1, 2, 3, 7, 10, and 11) were clustered at the N-terminus in all members of existed in all the members in the NHX family. Motif 6 was existed in Vac-and Endo-classes NHXs, three motifs (motif 5, 8, and 9) were only existed in Vac-class NHXs, and four motifs (motif 12, 13, 14, and 15) were only existed in PM-class NHXs. In addition, motif 4 was also Vac-class specific motif, but two AtNHXs (AtNHX3 and AtNHX4) in this subfamily were lost this motif (Fig. 4).

**Figure 4.**
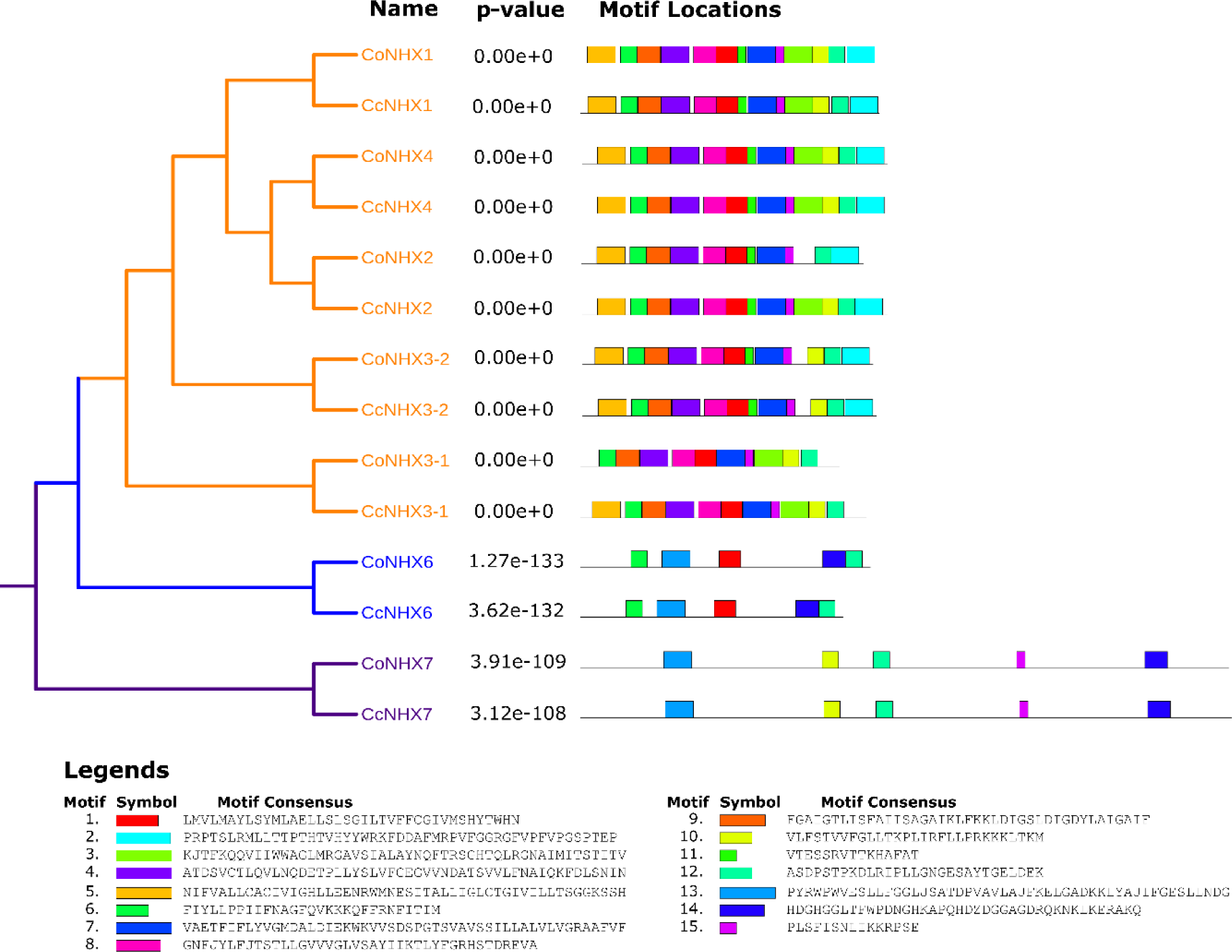
Motif analyses of Co and Cc NHXs. All motifs were identified by the MEME database with complete amino acid sequences of Co and Cc APQs. Length of motif for each AQPs protein are displayed proportionately. Details of the different motifs indicated by different colours are shown in supplementary Fig.1.

### Promoter profiling

Cis-elements played pivotal function in the regulation of gene expression by controlling the efficiency of the promoters. Studies on cis-elements could provide key foundation for further functional research of the NHXs gene family. In silico sequence analysis of promoter region showed presence of an important putative cis-acting element (Figure 5, Supplementary table 5-6), such as the ABA-response elements (ABREs) denoting possible ABA dependent regulation^56^, dehydration responsive elements (MBS DRE/CRT and G/ACCGCC), low temperature responsive elements (LTRE and CCGAC), heat shock elements (HSEs), cis-elements necessary for induction of many heat shock induced genes^57^, heat shock, osmotic stress, low pH, nutrient starvation responsible element (STRE), Auxin-responsive element (TGA), and hormone responsive elements (GARE motif, CGTCA-motif, TGACG-motif, and TCA).

**Figure 5.**
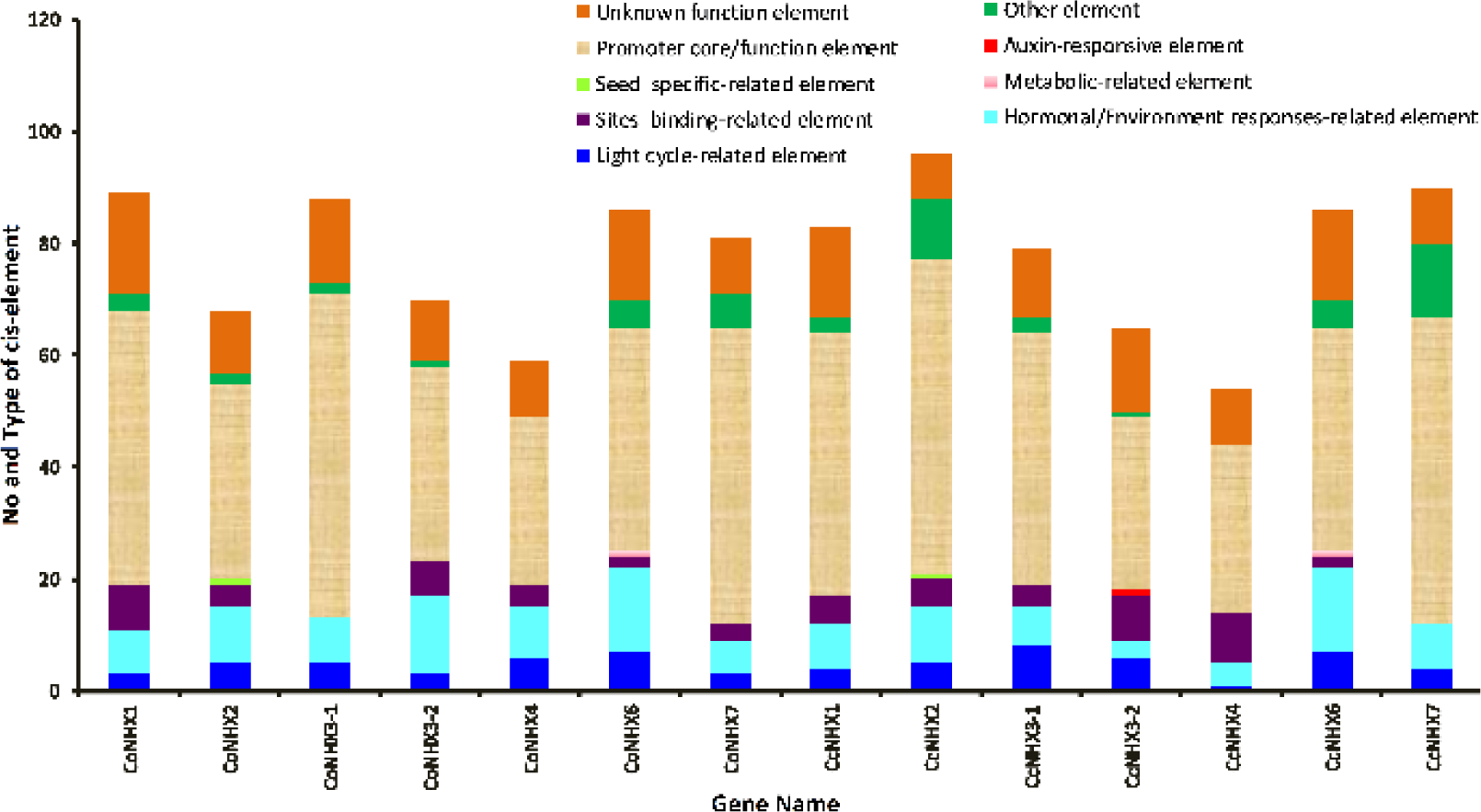
Promoter analysis of CoNHX and CcNHX genes. The numbers of different *cis*-elements were presented in the form of bar graphs; Different cis-elements with the same or similar functions are shown in the same color.

Presence of plant hormone responsive *cis-*elements such as AuxRRcore, ABRE, ERE, TCA-element, and GARE-motif in several Co and Cc NHXs, suggesting their expression regulation by different types of hormones. In this study PLACE database^43^ also found a series of cis-elements involved in abiotic stress responses, light and circadian rhythms regulation and seed development, Ca^2+^-response, Circadian regulation such as: ABRELATERD1, ACGTATERD1, ANAERO1CONSENSUS, ARR1AT, BIHD1OS, CAATBOX1, CACTFTPPCA1, CARGCW8GAT, CIACADIANLELHC, CCAATBOX1, CURECORECR, DOFCOREZM, DPBFCOREDCDC3, EBOXBNNAPA, GATABOX, GT1CONSENSUS, INRNTPSADB, MYBCORE, MYCCONSENSUSAT, POLASIG2, POLASIG3, POLLEN1LELAT52, SEF4MOTIFGM7S, TAAAGSTKST1, TATABOX2 and WRKY71OS. We hypothesize that external environmental stresses could induce the expression of Co and Cc NHX genes through their responsive cis-acting elements, further heightening the plant’s resistance to environmental stresses.

### Nomenclature of Corchorus species NHXs gene family

In accordance to the original nomenclature guidelines for plants genes, a two letter prefix derived from the genus and species names of the organisms in which the genes are present, for example At for Arabidopsis thaliana^58,59^. Co and Cc prefixes were used for Corchorus olitorius and Corchorus capsularis respectively where orthologs of phylogenetically related NHX genes of same clade with the model plant Arabidopsis and high percentage of similarity in amino acid sequence of paralogous genes (Supplementary Table 7) was the basis of classification and nomenclature^60^.

### Genomic distribution

Co and Cc NHX are obtained in cluster form in their respected genome. In C. olitorius 6 out of 7 were obtained and 5 out of 7 in C. capsularis) and 1 and 2 genes of C. olitorius and capsularis respectively were located on unanchored Scaffolds (Supplementary Table 8). According to supplementary table 4 Chromosome 2 and 5 contain two NHX genes genes each where as rest two genes are obtained in Chromosome 4 and 7. CoNHX5 obtained in unanchored scaffolds Scf7180001685718. In C. capsularis CcNHX1 and 3 are obtained in Chromosome 2 and CcNHX2, 4, 5 are found in Chromosome no 5,3 and 6. CcNHX6 and seven are in unanchored scaffold (scf7180000535309 and scf7180000535435).

### Expression profiles of the NHX genes under abiotic stress

Gene expression profile provides an important clue to determine gene functionality. The members of NHX gene family play important roles in response to various environmental stresses such as higher osmotic stress, high salinity and drought condition. The publicly available transcriptome dataset of jute against salinity and drought condition were used for NHX genes expression profiling (Fig.6 and supplementary table 9-10). In case of jute abiotic stress (Salinity and drought) transcriptome data showed that Co and Cc NHX1, 3, 6 and 7 are upregulated in drought condition in both species besides this CcNHX4 found that C. capsularis expression is higher in drought condition but C. olitorius showed lower expression. In case of salinity stressed transcriptome data, the Co and CcNHX2 and 6, showed higher expression in both jute species in both root and leaf data. On the other hand CcNHX7 showed upregulation C. capsularis and down regulation in C. olitorius. In addition, salinity stressed transcriptome the CcNHX1 expressed highly in the leaf and root system but in CoNHX1expressed high in leaf but lower expression in root in C. colitorius (Fig. 6 and supplementary table 9)

**Figure 6.**
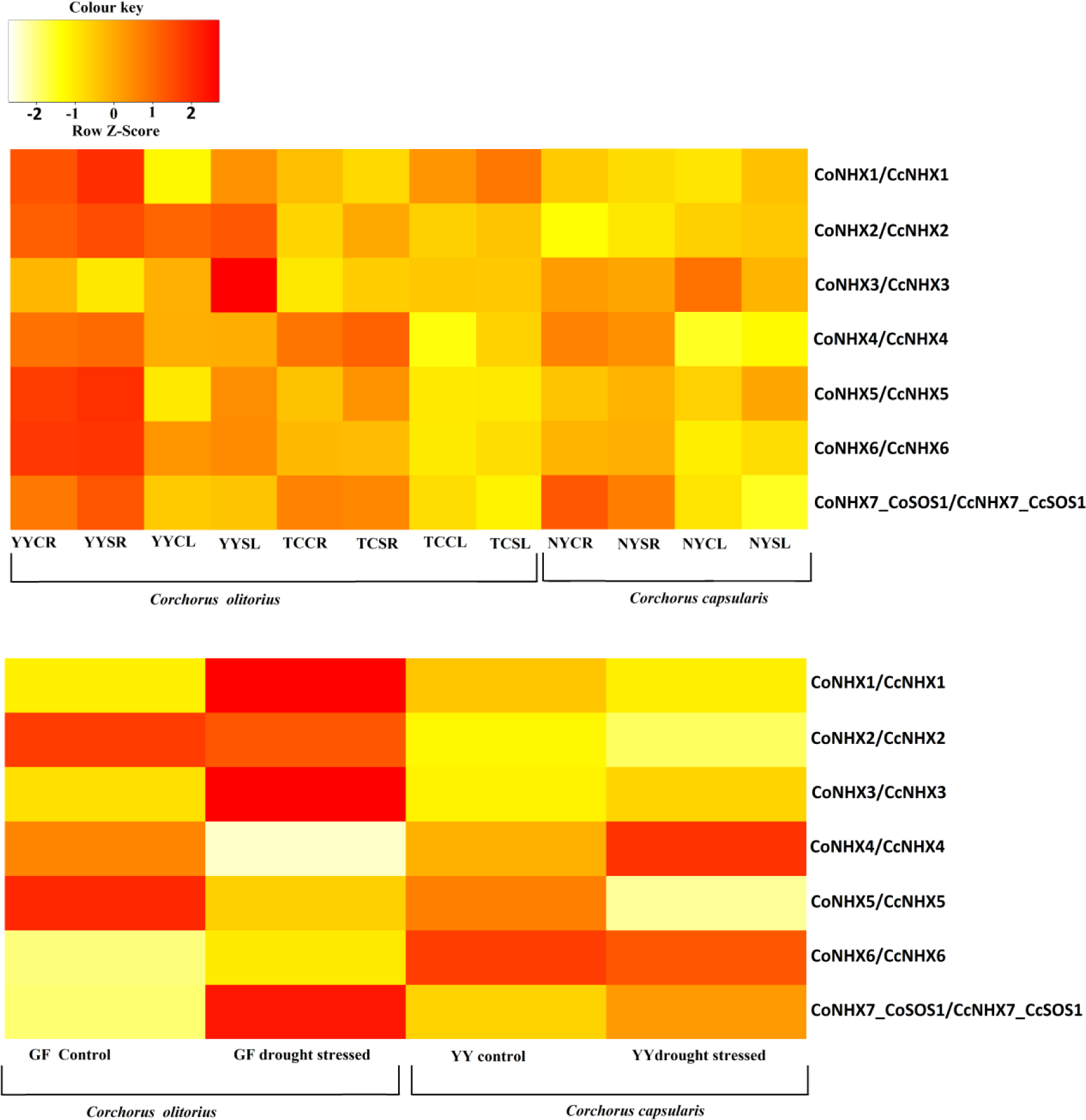
Expression analysis of Co and Cc NHXs responding to salinity and drought stress. YYCR= Capsularis variety YY control root; YYSR= Capsularis variety YY stress root; YYCL= Capsularis variety YY control leaf; YYSL = Capsularis variety YY Stress leaf; NYCR= Olitorius variety NY Control Root; NYSR = Olitorius variety NY Stress Root; NYCL= Olitorius variety NY Control Leaf; NYSL= Olitorius variety NY Stress leaf; TCCR= Olitorius saline tolerant TC accession Control Root; TCSR= Olitorius saline tolerant TC accession Stressed Root; TCCL = Olitorius saline tolerant TC accession Control Leaf; TCSL = Olitorius saline tolerant TC accession Stressed leaf.

### Expression patterns of both Corchorus species NHX genes under salinity stress

We have studied expression of Co and Cc NHX genes in the control and salinity treatment using qRT-PCR. The result of qRT-PCR analysis for the different treatments of the two Corchorus species NHX gene family members have very complex and different expression and induction patterns of NHX transcripts in leaf and root (Fig.7). NHX genes are highly expressed in leaf, and when both Corchorus species were exposed to salinity stress most of the genes were dramatically induced after each couple of hour interval. On the other hand the expression pattern in root system is lower than leaf (Fig.7). The orthologues of Co and Cc NHX7 and NHX1 was well known as salt tolerant locus similar to AtSOS1 and AtNHX1 in Arabidopsis, which plays a crucial role in root Na+ long distance response to leaf.

**Figure 7.**
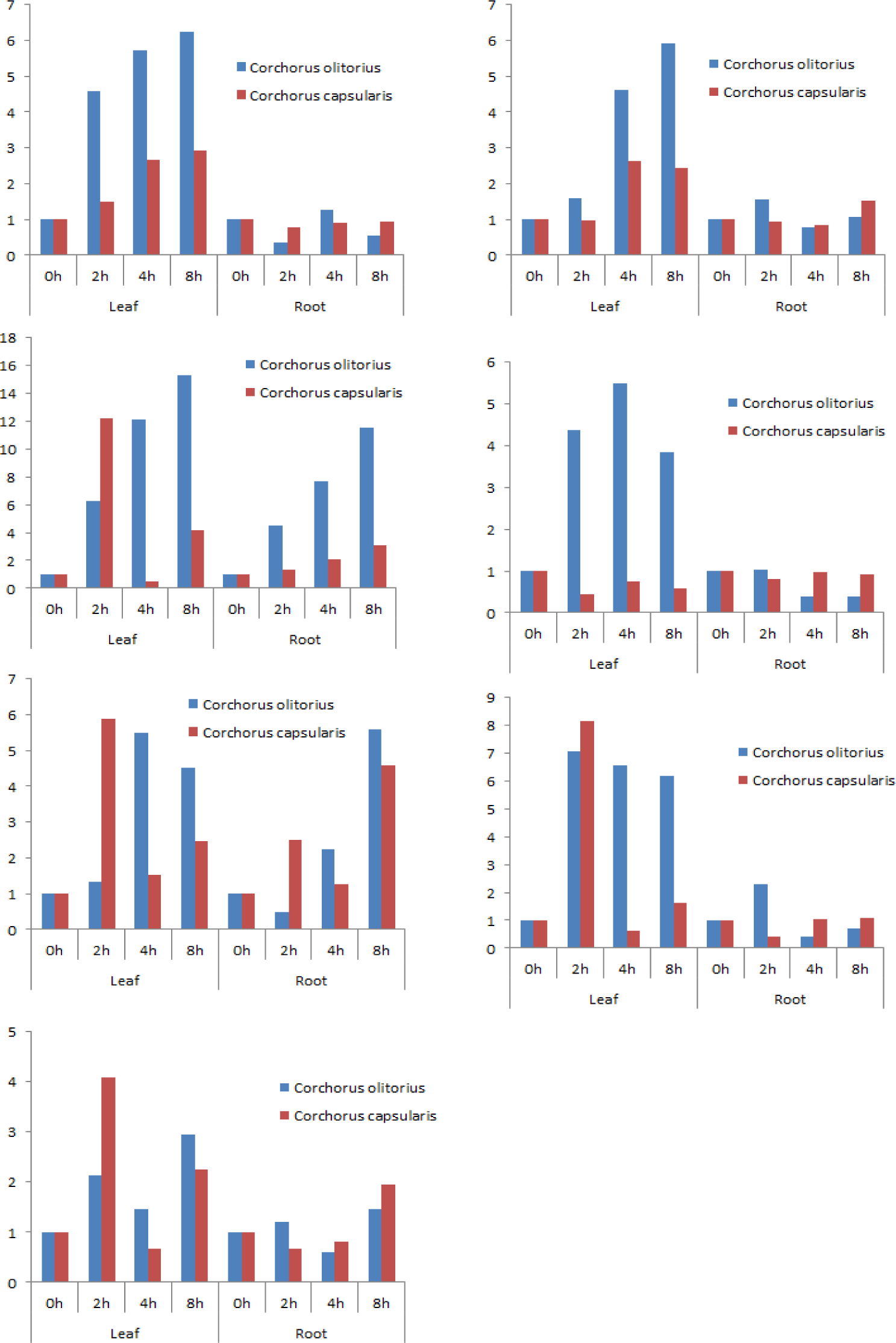
Expression profile of Co and CcNHXs in response to salt stress treatment. The qRT-PCR analysis were performed using total RNA extraction from root and leaftissue of C. olitorius cv O4 and C. capsularis cv CVL1 under high salinity stress (250mM NaCl for 0H, 2h, 4h, and 8h. The error bar represent the standard error of the mean of the three independent replicates.

## Discussion

Salinity is one of the most important abiotic stresses, affecting approximately 20% of the irrigated land used for agriculture globally^61^. To deal with salt stress, plants develop physiological, biochemical, and ecological strategies. Some common strategies include root ion uptake, root ion exclusion, ion accumulation in vacuoles of root or shoot cells, regulation of ion transport from root to shoot, increased tolerance for toxic ions, and compatible solute accumulation^1,62^. Although Cl^−^ toxicity is important in some sensitive plant species, Na^+^ toxicity is the primary aspect of the salinity stress ^63^. Traditionally, Na^+^ exclusion is believed to be the primary mechanism regulating salt tolerance^1,64^. However, recent findings indicate that only a fraction of the variation in salt tolerance is explained by Na^+^ exclusion, suggesting roles of other component traits in the salt tolerance mechanism^1,62,63,65,66^. It has been demonstrated that sequestration of Na^+^ into vacuoles is also an important strategy to keep the Na^+^ concentration low in cytosol^29,67^. Na^+^/H^+^ exchangers (NHXs) are the main players in sequestering Na^+^ into vacuoles and are shown to be involved in K^+^ homeostasis^29,36,68,69^. They are also involved in several cellular functions including cell expansion, stomatal regulation, pH control, vesicle trafficking, and flowering initiation^29^. Understanding the mechanism NHX family members in stress regulation and homeostasis is critical in comprehending the mechanisms regulating plant growth and development. Corresponding to eight NHX genes in Arabidopsis, seven were identified in both Corchorus species, of which five belonged to class I (vacuole specific, Co and CcNHX1-NHX5), while one belonged to class II (Golgi specific, Co and CcNHX6) and one to class III (plasma membrane specific, Co and CcNHX7) corroborated by phylogenetic analysis (Fig.2). Genes corresponding to AtNHX5 and AtNXH8 were missing in both Corchorus species. Lack of NHX5 in both Corchorus species may not be critical as NHX5 and NHX6 were shown to be functionally redundant; as a single mutant, nhx5 or nhx6 did not display any obvious phenotypes^22^. Additionally, either NHX5 or NHX6 was able to rescue the phenotype in the nhx5 nhx6 double mutant^22^. Finally, AtNHX8 is a Li^+^/H^+^ antiporter and plays role in Li^+^ extrusion but not for Na^+70^. Absence of NHX8 in both Corchorus species suggests differences in regulation of the Li^+^ detoxification mechanism.

Intron-exon structures also highlighted clear differences among three classes of Co and CcNHX genes with class I, class II, and class III genes containing 12-14, 18-21, and 23 exons, respectively (Fig. 3). Structural and domain analyses revealed that 12 TM domains and an amiloride binding motif are conserved in five Co and CcNHX Proteins (Fig. 1). The amiloride binding site is known to play an important role in inhibiting Na^+^/H^+^ exchange on amiloride binding^71^. However, TM5 and TM6 which are most critical for the antiport activity^72,73^ are conserved among all Co and CcNHX proteins (Fig.1). Based on the proposed ion transport mechanism for NHE1 in yeast, five residues (P167, P168, E262, D267, and S351) are known to play critical role in the antiport activity^74^. According to this model, P167 and P168 are predicated to lay the cation-transport path; E262 attracts H^+^ from the cytoplasmic side, which leads to protonation of D267, and S351 to exchange H^+^ with Na^+74,75^. Five corresponding residues, P91, P92, E183, D188, and S274, of Co and CcNHX1 are completely conserved in class I and class II (Fig. 1). AtNHX7/SOS1 is known to be a vital player in regulating Na^+^ efflux from roots^5^. It will be interesting to investigate if the SOS pathway-based regulation of Na^+^ exclusion from roots is maintained in both Corchorus species.

Like other NHX genes known in eukaryotes, the N-terminus of Co and CcNHX proteins is highly conserved as compared to the C-terminus^73^. Additionally, the C-termini of Co and CcNHX6 and NHX7 are diverged as compared to the class I (Fig. 1). C-terminus is proposed to be involved in differential regulation of the NHX genes, perhaps by posttranslational modifications such as phosphorylation, glycosylation or by facilitating protein-protein interactions^54,73^. During salt stress, when pH is high in the vacuole, Na^+^/H^+^ activity is favored and more Na^+^ is sequestered in to the vacuole^54^. It has been hypothesized that like SOS1, AtNHX1 may also be regulated through its interaction with SOS2^54^.

Expression analyses showed that all Co and CcNHX genes were upregulated in salinity treatment in leaf (Fig. 6). AtNHX3 is known to play an important role in K^+^/H^+^ exchange and facilitates more K^+^ accumulation in the vacuole as compared to Na^+76^. AtNHX6 is located in TGN, Golgi, and other endosomal compartments and plays a role in the trafficking of proteins to vacuoles^22^. NHX7/SOS1 is critical for excluding Na^+^ from plant roots^5^. Increased expressions of Co and CcNHX1, NHX2, NHX6, and NHX7 was observed in available transcriptome data and qRT-PCR support their roles in response to salt stress. Surprisingly, Co and CcNHX1 and NHX2, homologs of which helps in maintaining turgor regulation via active K^+^ uptake at the tonoplast during salt stress in Arabidopsis^67^, showed significant increment in gene expression in saline condition (Fig. 6). These observations suggest some functional similarities in NHX proteins among different plant species. There was no significant differences in expression profiles in leaves suggesting that in addition to sequestering Na^+^ into vacuoles, other component traits of the salt tolerance mechanism may be crucial in salinity tolerance. In-silico and expression analyses corroborates that NHX family members in both Corchorus species are involved in maintaining homeostasis in vacuoles and play important role in salinity stress.

## Conclusion

This is the first comprehensive genome-wide analysis of the NHX gene family in Corchorus species. Here we have identified and characterized seven Co and Cc NHX genes in each species and elucidated their structure, conserved motif, phylogenetic relation, cis-acting regulatory element and their potential involvement in terms of gene expression in response to abiotic stress. The global expression profiles of seven NHX genes obtained from C. olitorius and C. capsularis through RNA-seq analysis revealed higher expression of Co and CcNHX1, NHX2 NHX6 and NHX7 in drought and saline condition where as qRT-PCR analysis showed more upregulation in leaf than root against salinity, drought. Because of their importance in stress responses, NHX can be further investigated for the purpose of breeding and genetic improvement of jute in response to abiotic stress.

## Supporting information

Talbe S1-10

